# Age-accelerated cognitive decline in asymptomatic adults with CSF β-amyloid

**DOI:** 10.1101/220756

**Authors:** Lindsay R. Clark, Sara E. Berman, Derek Norton, Rebecca L. Koscik, Erin Jonaitis, Kaj Blennow, Barbara B. Bendlin, Sanjay Asthana, Sterling C. Johnson, Henrik Zetterberg, Cynthia M. Carlsson

**Affiliations:** Geriatric Research Education and Clinical Center, William. S. Middleton Memorial Veterans Hospital, Madison WI 53705, USA.; Alzheimer’s Disease Research Center, University of Wisconsin-Madison School of Medicine and Public Health, Madison, WI 53705, USA.; Wisconsin Alzheimer’s Institute, University of Wisconsin-Madison School of Medicine and Public Health, Madison, WI 53705, USA.; Medical Scientist and Neuroscience Training Programs, University of Wisconsin-Madison School of Medicine and Public Health, Madison, WI 53705, USA.; Department of Biostatistics and Medical Informatics, University of Wisconsin-Madison School of Medicine and Public Health, Madison, WI 53705, USA.; Clinical Neurochemistry Laboratory, Sahlgrenska University Hospital, Mölndal, Sweden.; Department of Psychiatry and Neurochemistry, Institute of Neuroscience & Physiology, The Sahlgrenska Academy at the University of Gothenburg, Molndal, Sweden; Department of Molecular Neuroscience, University College London, Institute of Neurology, Queen Square, London, UK

**Keywords:** Alzheimer’s disease, cognitive aging, cognitive neuropsychology in dementia, neuropsychological assessment, cerebrospinal fluid

## Abstract

**Objective:** Compare cognitive and hippocampal volume (HCV) trajectories in asymptomatic middle-aged and older adults with positive cerebrospinal fluid (CSF) markers of β-amyloid (Aβ) or tau to adults without an AD-associated biomarker profile.

**Method:** 392 adults enrolled in a longitudinal cohort study (Wisconsin Registry for Alzheimer’s Prevention or Wisconsin Alzheimer’s Disease Research Center) completed a lumbar puncture and at least two biennial or annual neuropsychological evaluations. Cutoffs for Aβ_42_, total tau, and phosphorylated tau were developed via receiver operating characteristic curve analyses on a sample of 78 participants (38 dementia, 40 controls). These cutoffs were applied to a separate sample of 314 cognitively healthy adults (mean age at CSF collection = 61.5) and mixed-effects regression analyses tested linear and quadratic interactions of biomarker group × age at each visit on cognitive and HCV outcomes.

**Results:** 215 participants (69%) were biomarker negative (preclinical AD Stage 0), 46 (15%) were Aβ+ only (preclinical AD Stage 1), 25 (8%) were Aβ+ and tau+ (preclinical AD Stage 2), and 28 (9%) were tau+ only. Both Stage 1 and Stage 2 groups exhibited greater rates of linear decline on story memory and processing speed measures, and non-linear decline on list-learning and set-shifting measures compared to Stage 0. The tau+ only group did not significantly differ from Stage 0 in rates of cognitive decline.

**Conclusion:** In an asymptomatic at-risk cohort, elevated CSF Aβ (with or without elevated tau) was associated with greater rates of cognitive decline, with the specific pattern of decline varying across cognitive measures.

## INTRODUCTION

Although most studies of preclinical Alzheimer’s disease (AD) focus on older adults, recent studies report that middle-aged adults with CSF biomarkers of both beta-amyloid (Aβ) and tau exhibit more rapid decline on cognitive and clinical measures than those with only one abnormal biomarker^1, 2^. These studies support guidelines defining preclinical AD as the presence of Aβ and neurodegeneration, while designating the presence of only one feature as “asymptomatic at-risk for AD” ^3^. However, prior studies examined change on cognitive composite scores or global screening measures and it remains unclear whether the presence of either Aβ or tau in isolation is associated with decline within specific cognitive domains, such as memory. Additionally, although cutoff values defining normal or abnormal levels of Aβ and tau are useful clinically, examining relationships between biomarkers and clinical symptoms along a continuum may provide additional information.

Our analysis was designed to replicate and build upon prior work by: 1) identifying Aβ and tau positivity in a longitudinal cohort sample of cognitively healthy middle-aged and older adults, 2) comparing biomarker groups on longitudinal neuropsychological performance across multiple measures and 3) investigating relationships between continuous variables of Aβ, tau, and cognitive performance. We hypothesized that adults with both Aβ and tau positivity would exhibit greater rates of cognitive decline compared to biomarker negatives. Based on prior work showing associations between Aβ and cognitive decline^4, 5^, we further hypothesized that those with Aβ+ would exhibit greater decline on memory measures, whereas tau+ adults would not differ from biomarker negatives.

## METHODS

### Participants

Participants included 392 middle-aged or older community-dwelling adults enrolled in longitudinal cohort studies of Wisconsin Registry for Alzheimer’s Prevention (WRAP)^6^ (*n*=141) or the Wisconsin Alzheimer’s Disease Research Center (WADRC) clinical core (*n*=251). These cohorts include cognitively healthy and impaired participants, are enriched for at-risk adults with family history of AD, and conduct study evaluations on an annual or biennial basis. Cognitive status was determined by consensus conference panel based on National Institute on Aging-Alzheimer’s Association (NIA-AA) criteria^7, 8^. The current study included participants with dementia in the development of CSF cutoff values, but included cognitively healthy middle-aged and older adults in all remaining analyses. Exclusion criteria consisted of only one study visit completed, relevant CSF or diagnosis data unavailable, diagnosis of mild cognitive impairment (MCI) or Impaired-not MCI at baseline or LP visit, or diagnosis of dementia that reverted to MCI at subsequent visits. Participants with incomplete neuropsychological data were included if data for at least two visits were available. Participants from the WRAP cohort were younger at baseline than those from the WADRC, but similar in sex distribution, education, and *APOE* genotype (see Table e-1).

### Standard Protocol Approvals, Registrations, and Patient Consents

The inclusion of human subjects in this study was approved by the University of Wisconsin-Madison Institutional Review Board and all participants provided informed consent.

### Procedures

CSF was collected in the morning after a minimum 12-hour fast. A Sprotte spinal needle was inserted into the L3-L4 or L4-L5 vertebral interspace and 22 mL of CSF was removed via gentle extraction into polypropylene syringes. Within 30 minutes of collection, the CSF was combined, gently mixed, centrifuged to remove red blood cells or other debris, aliquoted into 0.5-mL polypropylene tubes, and stored at −80°C. Samples were sent in batches at two time points for analysis at the Clinical Neurochemistry Laboratory at the Sahlgrenska Academy of the University of Gothenburg, Sweden. All samples were analyzed according to protocols approved by the Swedish Board of Accreditation and Conformity Assessment using one batch of reagents (intraassay coefficients of variation < 10%) for each batch. Board-certified laboratory technicians blinded to clinical diagnosis performed all analyses on one occasion for each of the two batches. CSF samples were assayed for total tau (t-tau), phosphorylated tau (p-tau_181_), amyloid beta 1–42 (Aβ42), and amyloid beta 1–40 (Aβ40) using commercially available enzyme-linked immunosorbent assay methods (INNOTEST assays, Fujirebio, Ghent, Belgium; Triplex assays, MSD Human Aβ peptide ultra-sensitive kit, Meso Scale Discovery, Gaithersburg, MD). Additional details on batch-to-batch conversions is provided in the Supplemental Material.

A comprehensive neuropsychological assessment was completed at each visit. Measures of memory (Rey Auditory Verbal Learning Test [RAVLT] Total Trials 1-5 and Delayed Recall ^9^, Wechsler Memory Scale-Revised Logical Memory Story A [LM] Immediate and Delayed Recall ^10^) and executive functioning (Trailmaking Test Part B [TMT-B] ^11^, Animal Fluency, Wechsler Adult Intelligence Scale-Revised Digit Symbol ^12^) were included based on prior meta-analyses indicating that these cognitive domains demonstrate significant decline and associations with AD biomarkers in preclinical AD ^13-15^. A subset of 205 participants completed at least two MRI scans and were included in secondary analyses of hippocampal volume (HCV) change (see Supplemental Material for MRI details).

### Statistical analyses

Statistical analyses were conducted in R version 3.3.1 ^16^. Cutoff values for CSF assays were developed using receiver operating characteristic (ROC) curve analysis in the pROC package (version 1.8) ^17^ in 38 participants with clinical diagnoses of dementia due to AD based on NIA-AA criteria^8^ without reference to CSF biomarkers and 40 late middle-age (ages 48-64) stable cognitively healthy adults at lower risk for AD (*APOE* ε4 non-carrier, no family history of AD). Youden’s J (sensitivity + specificity −1), which maximizes both the sensitivity and specificity of a diagnostic test, was used.

To reduce potential risk of researcher assessment bias, a non-overlapping sample of 314 cognitively healthy participants (mean LP age 61.5) were included in subsequent analyses. We compared biomarker groups on demographic characteristics using chi-square and analysis of variance (ANOVA) tests. We compared mean neuropsychological performance and HCV among biomarker groups at the visit closest to the LP using analysis of covariance (ANCOVA) models with age at LP (mean = 61.5), sex (reference group = female), and years of education (mean = 16.3) as covariates. Comparisons of HCV also included TIV (mean = 1464.8 mm^3^) as a covariate.

To test if longitudinal change on the seven neuropsychological measures and HCV varied across biomarker groups, linear mixed-effects models were conducted using the lme4 package version 1.1-12^18^. Fixed effects included sex, years of education, practice effects (number of exposures to test ^19^), biomarker group (4 levels), age (at each visit), and the interaction of age × biomarker group. To allow for acceleration of cognitive decline with increasing age, two quadratic terms, age^2^ and age^2^ × biomarker group, were included in all models and removed if non-significant. To minimize collinearity in the linear and quadratic age terms, the age variable was centered on the sample mean. All models included random effects of intercept and slope nested within subject. The overall significance of the interaction term was assessed by likelihood ratio tests comparing the primary model and a model that did not include the interaction term. *P*-values for fixed effect coefficients were calculated using asymptotic properties of the estimates^20^. Statistical significance was defined as *p* < .05.

To investigate the relationship between cognitive or HCV change and continuous Aβ42 or tau values, we conducted two identical models to those above (excluding biomarker group terms). The first included predictors of Aβ42 (centered), p-tau (centered), age × Aβ42, age × p-tau, Aβ42 × p-tau, and age × Aβ42 × p-tau. The second model included effects of p-tau/Aβ42 and age × p-tau/Aβ42. Since t-tau was highly correlated with p-tau (*r* = .85, *p* < .001), we only included p-tau in these models.

## RESULTS

### Biomarker cutoffs

Table 1 details sample characteristics. All biomarker cutoffs had a minimum sensitivity and specificity of 70% and 90%, respectively (Table e-2). The ratios of tau to Aβ42 exhibited sensitivities and specificities ≥ 90% and greater area under the curve (AUC) values than Aβ42 (*p* < .05), Aβ42/Aβ40 (*p* < .05), and p-tau (*p* < .01).

**Table 1.** Sample characteristics

| Variable | ROC sample | Cognitively Healthy sample |
| --- | --- | --- |
| *N* (Total = 392) | 78<sup>a</sup> | 314 |
| Age at visit 1 (mean, range) | 64.5 (47-92) | 58.8 (37-85) |
| Age at lumbar puncture (LP) visit (mean, range) | 65.4 (48-93) | 61.5 (43-86) |
| Months between visit 1 and LP (mean, range) | 10.9 (0-91) | 33.5 (0-134) |
| Female (*n*, %) | 42 (54%) | 218 (69%) |
| Education (mean, SD, range) | 15.3 (2.6; 8-20) | 16.3 (2.5; 8-25) |
| *APOE* ε4+ (*n*, %) | 27 (35%) | 135 (43%) |
| Years in study (mean, SD, range) | 2.8 (2.6; 0-11) | 5.8 (3.5; 1-13) |
| Natural-log beta-amyloid ($A\beta_{42}$) (mean, SD, range) | 6.3 (0.4; 5.3-7.2) | 6.5 (0.3; 5.6-7.5) |
| Total tau (t-tau) (mean, SD, range) | 528.1 (366.0; 67.2-1633.0) | 324.7 (153.3; 67.2-1085.0) |
| Phosphorylated tau (p-tau) (mean, SD, range) | 58.2 (29.6; 17.1-152.0) | 43.5 (15.5; 12-114) |
| $A\beta_{42}/A\beta_{40}$ (mean, SD, range) | 0.08 (0.03; 0.04-0.13) | 0.1 (0.02; 0.04-0.2) |
| t-tau/$A\beta_{42}$ (mean, SD, range) | 1.2 (1.1; 0.1-4.5) | 0.5 (0.4; 0.1-3.5) |
| p-tau/$A\beta_{42}$ (mean, SD, range) | 0.1 (0.1; 0.03-0.5) | 0.1 (0.04; 0.02-0.3) |
| Diabetes | 6 (8%) | 16 (5%) |
| Hypertension | 28 (36%) | 61 (19%) |
| Hypercholesterolemia | 34 (44%) | 122 (39%) |
| History of stroke or TIA | 3 (4%) | 0 (0%) |
| Prescribed cognitive-enhancing medication<sup>b</sup> | 37 (47%) | 0 (0%) |

ROC=Receiver Operator Characteristic Curve; AD=Alzheimer's disease; *APOE*=Apolipoprotein
<sup>a</sup>*n*=40 cognitively healthy controls (51%) and *n*=38 participants with clinical diagnoses of dementia due to Alzheimer's disease (49%); <sup>b</sup>Donepezil, Memantine, Galantamine, or Rivastigmine

### Characteristics of biomarker groups

Of the 314 cognitively healthy participants, 53 (17%) had a positive tau biomarker (either p-tau ≥ 59.5 (*n*=40; 13%) or t-tau ≥ 461.26 (*n*=42; 13%)) and 76 (24%) had a positive amyloid biomarker (either Aβ42(ln) ≤ 6.156 [back-transformed value = 471.54] (*n*=44; 14%) or Aβ42/Aβ40 ≤ 0.09 (*n*=67; 21%)).

The majority of participants were negative for both biomarkers of Aβ and tau (Stage 0 = 68.5%). 14.6% were positive for Aβ only (Stage 1), 8% were positive for tau only, and 8.9% were positive for both Aβ and tau (Stage 2). Stages 0 and 1 did not differ on mean t-tau (*p* = .10) or p-tau (*p* = .41). Stage 0 had lower Aβ42 than the tau+ group (*p* < .001), but did not differ on the Aβ42/Aβ40 ratio (*p* = .97). The Stage 2 group was the oldest and the Stage 0 group was the youngest (*p* < .001). The Stage 1 and 2 groups included greater proportions of *APOE* ε4 carriers (63 and 64% respectively) compared with the Stage 0 or tau+ groups (28 and 38%). There were no differences between biomarker groups in sex, years of education, family history of AD, or source cohort (Table 2).

**Table 2.**
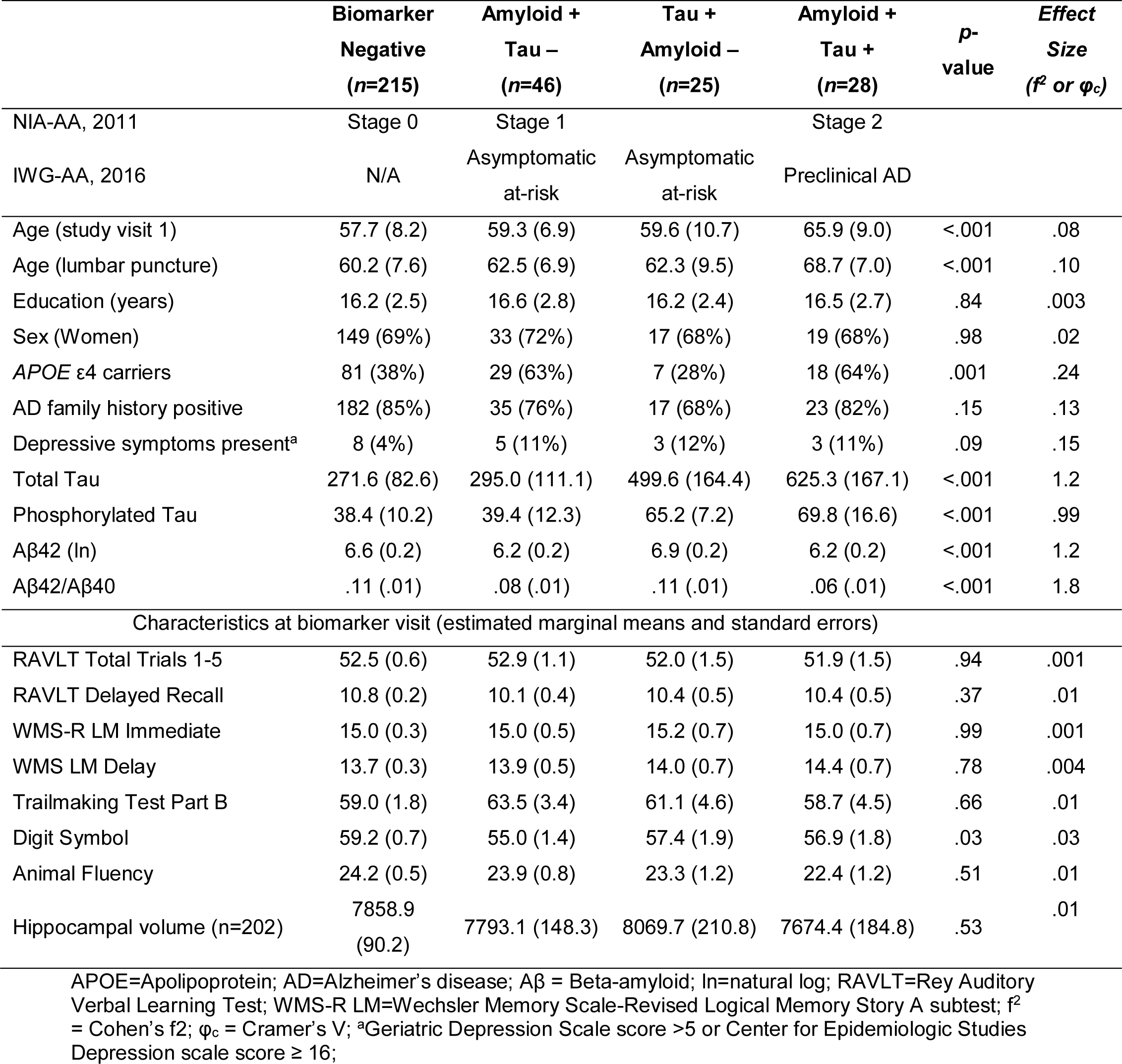
Biomarker group characteristics (*n*=314) (mean [SD] or n (%))

|  | **Biomarker**<br>**Negative**<br>**($n=215$)** | **Amyloid +**<br>**Tau –**<br>**($n=46$)** | **Tau +**<br>**Amyloid –**<br>**($n=25$)** | **Amyloid +**<br>**Tau +**<br>**($n=28$)** | **$p$-**<br>**value** | **Effect**<br>**Size**<br>**($f^2$ or $\phi_c$)** |
| --- | --- | --- | --- | --- | --- | --- |
| NIA-AA, 2011 | Stage 0 | Stage 1 |  | Stage 2 |  |  |
| IWG-AA, 2016 | N/A | Asymptomatic<br>at-risk | Asymptomatic<br>at-risk | Preclinical AD |  |  |
| Age (study visit 1) | 57.7 (8.2) | 59.3 (6.9) | 59.6 (10.7) | 65.9 (9.0) | <.001 | .08 |
| Age (lumbar puncture) | 60.2 (7.6) | 62.5 (6.9) | 62.3 (9.5) | 68.7 (7.0) | <.001 | .10 |
| Education (years) | 16.2 (2.5) | 16.6 (2.8) | 16.2 (2.4) | 16.5 (2.7) | .84 | .003 |
| Sex (Women) | 149 (69%) | 33 (72%) | 17 (68%) | 19 (68%) | .98 | .02 |
| APOE $\epsilon 4$ carriers | 81 (38%) | 29 (63%) | 7 (28%) | 18 (64%) | .001 | .24 |
| AD family history positive | 182 (85%) | 35 (76%) | 17 (68%) | 23 (82%) | .15 | .13 |
| Depressive symptoms present<sup>a</sup> | 8 (4%) | 5 (11%) | 3 (12%) | 3 (11%) | .09 | .15 |
| Total Tau | 271.6 (82.6) | 295.0 (111.1) | 499.6 (164.4) | 625.3 (167.1) | <.001 | 1.2 |
| Phosphorylated Tau | 38.4 (10.2) | 39.4 (12.3) | 65.2 (7.2) | 69.8 (16.6) | <.001 | .99 |
| Aβ42 (ln) | 6.6 (0.2) | 6.2 (0.2) | 6.9 (0.2) | 6.2 (0.2) | <.001 | 1.2 |
| Aβ42/Aβ40 | .11 (.01) | .08 (.01) | .11 (.01) | .06 (.01) | <.001 | 1.8 |
| Characteristics at biomarker visit (estimated marginal means and standard errors) |  |  |  |  |  |  |
| RAVLT Total Trials 1-5 | 52.5 (0.6) | 52.9 (1.1) | 52.0 (1.5) | 51.9 (1.5) | .94 | .001 |
| RAVLT Delayed Recall | 10.8 (0.2) | 10.1 (0.4) | 10.4 (0.5) | 10.4 (0.5) | .37 | .01 |
| WMS-R LM Immediate | 15.0 (0.3) | 15.0 (0.5) | 15.2 (0.7) | 15.0 (0.7) | .99 | .001 |
| WMS LM Delay | 13.7 (0.3) | 13.9 (0.5) | 14.0 (0.7) | 14.4 (0.7) | .78 | .004 |
| Trailmaking Test Part B | 59.0 (1.8) | 63.5 (3.4) | 61.1 (4.6) | 58.7 (4.5) | .66 | .01 |
| Digit Symbol | 59.2 (0.7) | 55.0 (1.4) | 57.4 (1.9) | 56.9 (1.8) | .03 | .03 |
| Animal Fluency | 24.2 (0.5) | 23.9 (0.8) | 23.3 (1.2) | 22.4 (1.2) | .51 | .01 |
| Hippocampal volume ($n=202$) | 7858.9<br>(90.2) | 7793.1 (148.3) | 8069.7 (210.8) | 7674.4 (184.8) | .53 | .01 |

APOE=Apolipoprotein; AD=Alzheimer's disease; Aβ = Beta-amyloid; ln=natural log; RAVLT=Rey Auditory Verbal Learning Test; WMS-R LM=Wechsler Memory Scale-Revised Logical Memory Story A subtest; $f^2$ = Cohen's $f^2$ ; $\phi_c$ = Cramer's V; <sup>a</sup>Geriatric Depression Scale score >5 or Center for Epidemiologic Studies Depression scale score $\geq 16$ ;

### Cognitive trajectories across biomarker groups

At the visit closest to the LP, there were no significant differences in cognitive performance or HCV across biomarker groups (Table 2), with the exception of processing speed (Digit Symbol).

Longitudinal neuropsychological performance for each biomarker group is displayed in Figure 1. Results from likelihood ratio tests (*χ*^2^(3)) indicated that age^2^ × biomarker group accounted for a significant amount of variation in change on RAVLT Delay (*χ*^2^ = 9.74, *p* = .02) and similar but non-significant variation in change on RAVLT Total (*χ*^2^ = 7.11, *p* = .07) and TMT-B (*χ*^2^ = 6.89, *p* = .08). Compared to the Stage 0 group, both Stage 1 and 2 groups showed more rapid, non-linear decline with age on the RAVLT Delay (*p’s* < .05), whereas the Stage 2 group only showed more rapid, non-linear decline on the RAVLT Total (*p* = .02). Compared to the Stage 0 group, the Stage 1 group showed more rapid, non-linear change with age on TMT-B (*p* = .02). In contrast, the tau+ group did not significantly differ from the Stage 0 group. Age^2^ × biomarker group was non-significant for the remaining outcomes (*p’s* > .43). Model parameters are displayed in Table 3.

**Figure 1.**
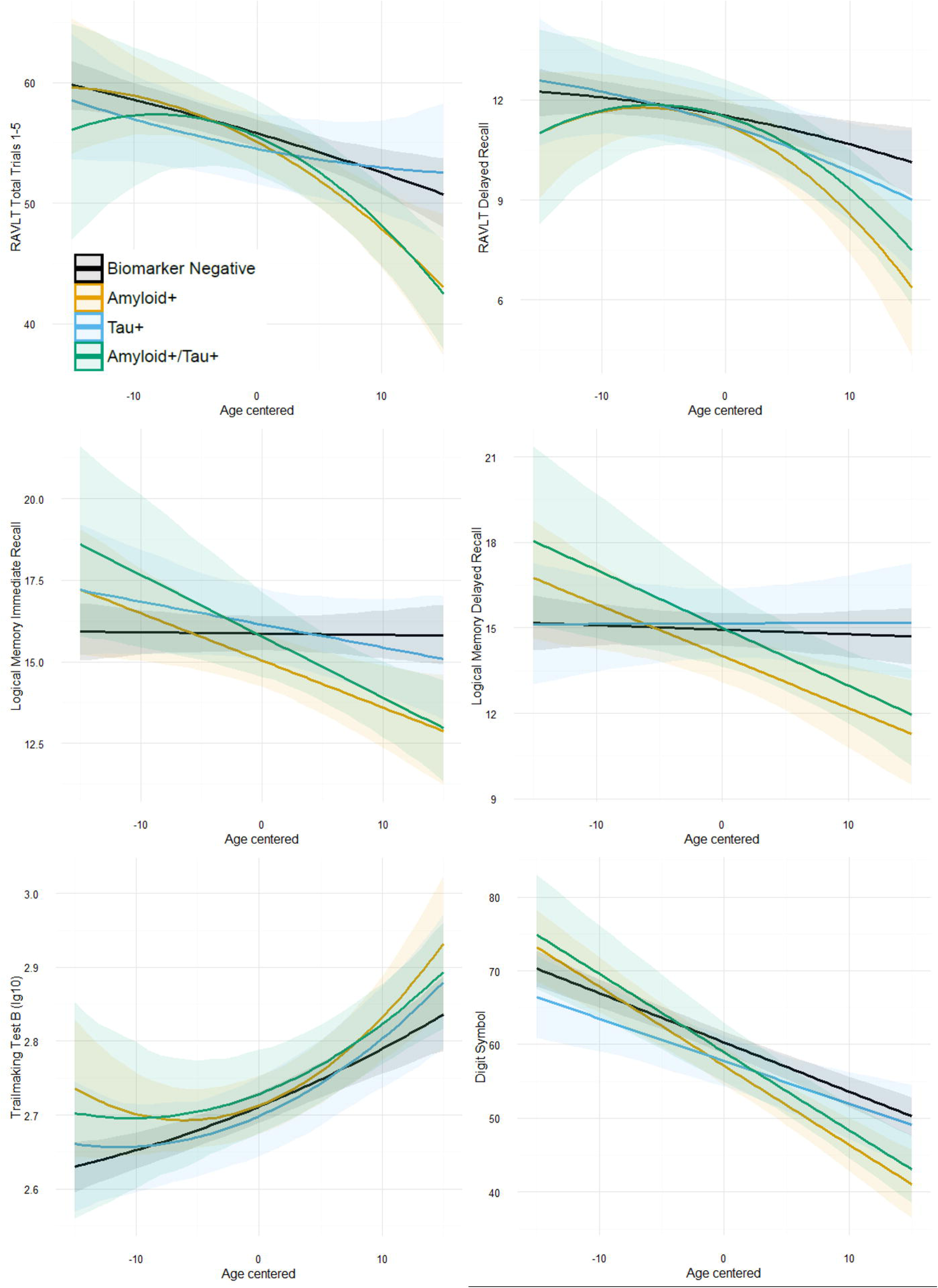
Biomarker groups and cognitive trajectories. Graphs depict neuropsychological performance on the y-axis for six cognitive measures and age at each visit (centered on mean age) on the x-axis. Each line depicts the estimated slope for the four biomarker groups, adjusting for covariates of sex, education, and practice effects. Higher scores equate better performance on all measures except TMT-B (higher scores = worse performance). Quadratic terms were retained for the RAVLT and TMT-B. Non-significant quadratic terms were removed for other outcomes and linear effects are depicted. Both the Aβ+ only group (orange) and the Aβ+/Tau+ group (green) exhibited significantly greater decline than the biomarker negative group (black). In contrast, the group with only tau+ (blue) did not differ from biomarker negative individuals.

**Table 3.** Parameter estimates from linear mixed-effects models

|  | RAVLT<br>TOTAL<br>TRIALS 1-5 | RAVLT<br>DELAYED<br>RECALL | LM<br>IMMEDIATE<br>RECALL | LM<br>DELAYED<br>RECALL | DIGIT<br>SYMBOL | TRAILS B<br>(lg10) |
| --- | --- | --- | --- | --- | --- | --- |
| Fixed Effects | B (SE) | B (SE) | B (SE) | B (SE) | B (SE) | B (SE) |
| Intercept | 45.0 (2.5)*** | 8.8 (0.9)*** | 9.4 (1.0)*** | 8.3 (1.1)*** | 50.2 (3.2)*** | 2.8 (0.0)*** |
| Biomarker Group |  |  |  |  |  |  |
| Aβ-/Tau- | -- | -- | -- | -- | -- | -- |
| Aβ+ | -0.7 (1.2) | -0.3 (0.4) | -0.8 (0.4) | -0.9 (0.5) | -3.2 (1.4)* | 0.001 (0.0) |
| Tau+ | -1.3 (1.6) | -0.3 (0.5) | 0.3 (0.6) | 0.2 (0.6) | -2.5 (1.8) | -0.01 (0.0) |
| Aβ+/Tau+ | -0.3 (1.6) | -0.02 (0.6) | -0.1 (0.7) | 0.07 (0.7) | -1.3 (2.0) | 0.02 (0.0) |
| Age each visit (center) | -0.3 (0.1)*** | -0.1 (0.0)*** | -0.004 (0.0) | -0.02 (0.0) | -0.7 (0.1)*** | 0.01 (0.0)*** |
| Age each visit (center)<sup>2</sup> | -0.002 (0.0) | -0.002 (0.0) | -- | -- | -- | 0.0001 (0.0) |
| Sex (male) | -6.8 (0.8)*** | -1.8 (0.3)*** | -2.1 (0.3)*** | -2.1 (0.4)*** | -3.4 (1.1)** | 0.03 (0.0)* |
| Education (years) | 0.4 (0.2)** | 0.1 (0.1)* | 0.4 (0.1)*** | 0.3 (0.1)*** | 0.5 (0.2)* | -0.004 (0.0) |
| Practice Effect | 1.2 (0.1)*** | 0.2 (0.0)*** | 0.2 (0.1)** | 0.3 (0.1)*** | 0.8 (0.2)*** | -0.01 (0.0)*** |
| Age each visit x Group |  |  |  |  |  |  |
| Age x Aβ-/Tau- | -- | -- | -- | -- | -- | -- |
| Age x Aβ+ | -0.2 (0.1)* | -0.1 (0.0)* | -0.1 (0.1)* | -0.2 (0.1)** | -0.4 (0.1)* | -.0003 (0.0) |
| Age x Tau+ | 0.1 (0.1) | -0.05 (0.0) | -0.1 (0.1) | 0.02 (0.1) | 0.1 (0.2) | .0004 (0.0) |
| Age x Aβ +/Tau+ | -0.2 (0.2) | -0.05 (0.1) | -0.2 (0.1)** | -0.2 (0.1)** | -0.4 (0.2)* | -.0005 (0.0) |
| Age each visit<sup>2</sup> x Group |  |  |  |  |  |  |
| Age<sup>2</sup> x Aβ-/Tau- | -- | -- | -- | -- | -- | -- |
| Age<sup>2</sup> x Aβ+ | -0.01 (0.0) | -0.01 (0.0)* | -- | -- | -- | .0004 (0.0)* |
| Age<sup>2</sup> x Tau+ | 0.01 (0.0) | -0.001 (0.0) | -- | -- | -- | .0002 (0.0) |
| Age<sup>2</sup> x Aβ +/Tau+ | -0.03 (0.0)* | -0.01 (0.0)* | -- | -- | -- | .0002 (0.0) |

RAVLT=Rey Auditory Verbal Learning Test; WMS-R LM=Wechsler Memory Scale-Revised Logical Memory Story A subtest; \*\*\*p ≤ .001; \*\*p ≤ .01; \*p ≤ .05
Note: Quadratic terms were non-significant for LM and Digit Symbol measures. Final models with quadratic terms removed are reported here.

For the remaining outcomes (in which the quadratic term was not associated with cognitive performance), results from likelihood ratio tests (*χ*^2^(3)) indicated that the interaction between age × biomarker group accounted for a significant amount of variation in change on LM Immediate (*χ*^2^ = 11.74, *p* < .01), LM Delay (*χ*^2^= 12.77, *p* < .01), and Digit Symbol (*χ*^2^ = 13.21, *p* < .01). For all three outcomes, Stage 1 and Stage 2 exhibited greater age-related decline than Stage 0 (*p* < .05). In contrast, the tau+ group did not differ from the Stage 0 group in rates of cognitive change. Age-related change in HCV did not differ by biomarker group. Sensitivity analyses conducted on WRAP and ADRC cohorts separately revealed similar directions of effects, but slight heterogeneity in magnitude of beta-weights possibly due to baseline age differences across cohorts (see Supplemental Material).

### Cognitive trajectories and continuous CSF values

There were no significant interactions between Aβ42 × p-tau × age^2^; this term was removed from subsequent analyses. The three-way interaction between Aβ42 × p-tau × age was statistically significant for LM Delay (*B* = .01, *p* = .03), in which the relationship between p-tau and longitudinal story memory performance was dependent on Aβ42. Similar to results above, two-way interactions between age^2^ × Aβ42 were significant for RAVLT Delay (age^2^: *B =* -.004, *p* < .01; age^2^ × Aβ42: *B =* 0.01, *p*=.03), RAVLT Total (age^2^: *B =* -.01, *p* = .05; age^2^ × Aβ42: *B* = 0.03, *p* < .01), and TMT-B (age^2^: *B* = .0002, *p* < .001; age^2^ × Aβ42: *B* = -.0004, *p* = .03), indicating that lower CSF Aβ42 (higher brain amyloid) was associated with greater non-linear decline. For outcomes for which age^2^ × Aβ42 was non-significant, greater amyloid burden was associated with greater linear decline (significant age × Aβ42 interaction) for LM Immediate (*B* = .22, *p* = .001), LM Delay (*B =* 0.22, *p* < .01), Digit Symbol (*B* = .48, *p* < .01), and Animal Fluency (*B* = .34, *p* < .01). In contrast, there were no interactions between age (linear or quadratic) × p-tau (Figure 2).

**Figure 2.**
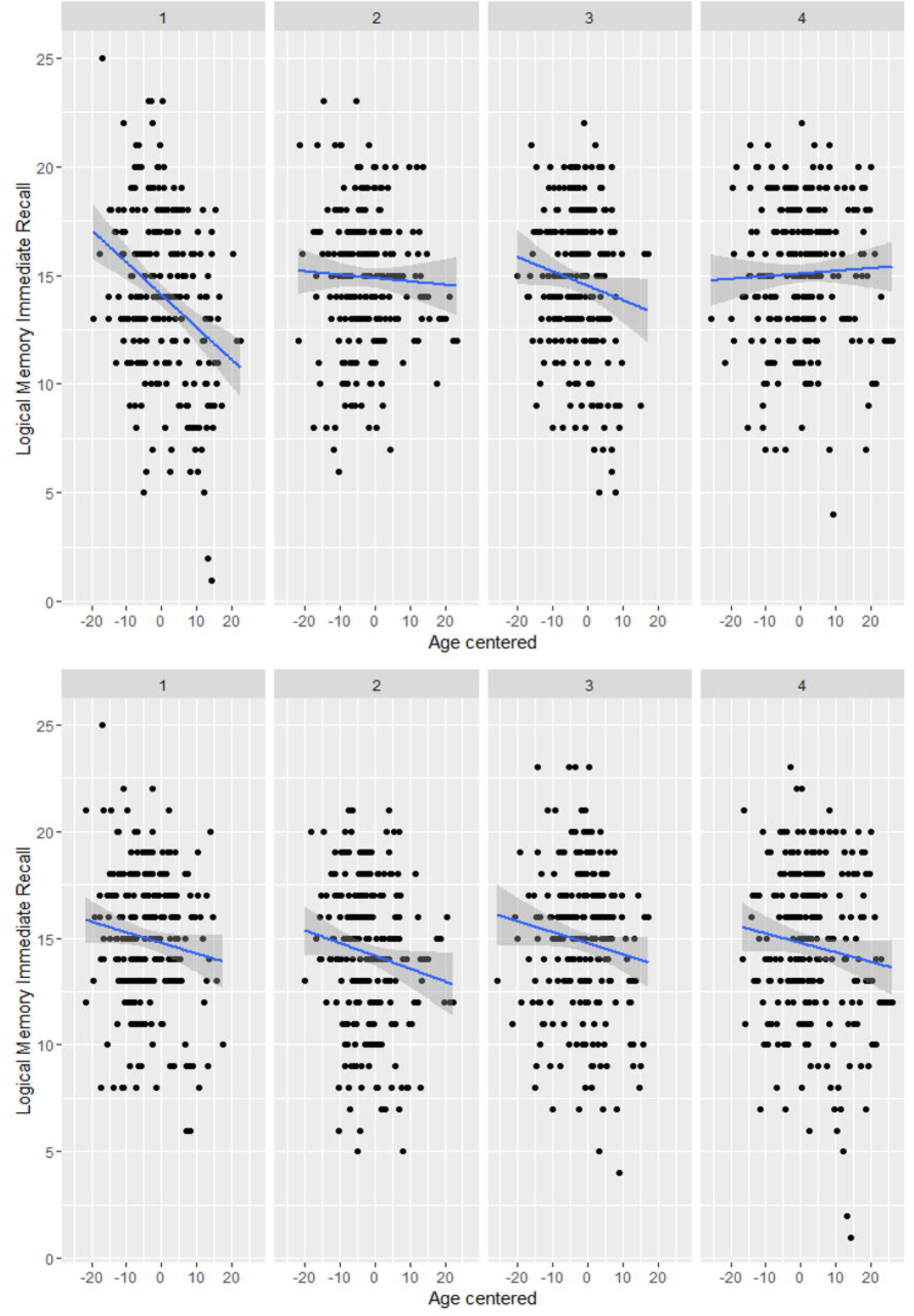
Relationships between Aβ42, tau, and longitudinal verbal memory performance. Two-way interaction between age at each visit and Aβ42 (top) or ptau (bottom) on memory performance. Figures depict that although performance generally decreases with age, those with low Aβ42 (high brain amyloid) exhibit most rapid decline, whereas the association between age at each visit and memory performance does not vary by ptau. Facets depict biomarker level by quartile (1 = lowest quartile (0-25%), 2= 25-50%, 3 = 50-75%, 4 = highest quartile (75-100%)).

Age^2^ × p-tau/Aβ42 was significant for TMT-B (*B* = .004, *p* < .01) and marginal for RAVLT Delay (*B* = −0.1, *p* = .08). Age × p-tau/Aβ42 was significant for all other outcomes with the exception of HCV, indicating that elevated AD biomarkers were associated with greater decline on RAVLT Total (*B* = −3.6, *p* < .01), LM Immediate (*B* = −2.3, *p* < .001), LM Delay (*B* = −2.3, *p* < .001), Digit Symbol (*B* = −3.0, *p* = .02), and Animal Fluency (*B* = −2.6, *p* < .01).

## DISCUSSION

In 314 cognitively healthy middle-aged and older adults enriched for AD risk, approximately one-third were positive for CSF biomarkers of AD (Aβ or tau). Those with Aβ positivity (with or without tau positivity) exhibited significantly greater decline on neuropsychological measures than biomarker negative adults, whereas those with only tau positivity did not differ from biomarker negatives.

These results have potentially important implications pertaining to AD during the asymptomatic or preclinical period. First, 24% of the sample were Aβ positive and 17% were tau positive using the selected biomarker threshold at relatively young ages of 59.3 and 59.6 for the Aβ only and tau only groups respectively, and 65.9 for the Aβ and tau positive group. While the age of the latter group was significantly older than other groups, the ages were overall quite young and empirically support the hypothesis^21^ that AD neuropathology changes begin well in advance of MCI and dementia syndromes.

Second, elevated Aβ in the absence of tau was associated with cognitive decline in late middle-age. This is an important finding because it adds to the debate on whether Aβ or tau more strongly contribute to early symptoms of cognitive decline. Although emerging evidence indicates that elevated Aβ on a PET scan is associated with increased risk for cognitive decline^4, 5, 22^, simultaneous measures of tau have not always been available, and therefore it is unclear whether results from prior studies are due to elevated Aβ alone or elevated Aβ and tau. Neuropathology studies demonstrating correlations between patterns of cognitive impairment in older adults with dementia and regional distribution of neurofibrillary tangle development^23, 24^ suggest that tau distribution drives major cognitive symptoms. However, the current results suggest that elevated Aβ independent of tau in late middle-age is associated with cognitive decline. Decline in this context was significant, but mild (e.g., using our regression results we estimate that 5-year decline on the RAVLT Total from age 61.5 to age 66.5 for the Aβ only group would be 3.2 points compared to 1.6 points for the biomarker negative group), and few individuals declined to a cognitively impaired diagnosis during the visits included in this study (e.g., only 4 participants declined from cognitively normal to MCI at the most recent visit). This finding in the context of the literature suggests that Aβ may be associated with subtle decline in midlife, whereas tau may contribute to more pronounced clinical symptoms as the disease progresses.

Third, the pattern of decline with age and Aβ varied across cognitive measures. Prior investigations of preclinical biomarker stage and longitudinal cognition in late middle-age have examined change on a global cognitive screener^2^ or composite score^1^; current results suggest examination of multiple cognitive domains may be useful in parsing out subtle patterns of decline related to Aβ. Specifically, performance on story memory and processing speed measures declined linearly with age and Aβ burden, whereas nonlinear decline on list-learning and set-shifting tasks indicated faster rates of decline on these measures with advancing age in the presence of Aβ burden. These results have potentially important implications for choosing appropriate outcome measures in clinical trials. For example, if a trial is enrolling older adults, it may be more optimal to choose a list-learning memory measure since it would be expected to decline more rapidly in older adults with AD pathology. Moreover, our results suggest that a neuropsychological measure of processing speed and working memory (Digit Symbol) may be a very early predictor of decline as this was the only cognitive measure that distinguished biomarker groups cross-sectionally at the biomarker visit. This is consistent with a prior study in a separate middle-aged cohort which reported that baseline performance on Digit Symbol and 3 additional measures best predicted conversion from cognitively normal to cognitively impaired^25^. Lastly, results across the majority of models including continuous CSF markers were similar to those using a group variable based on cutoffs (e.g., lower CSF Aβ42 was associated with worsening performance, whereas elevated tau was not). This finding suggests that dichotomizing continuous biomarker variables does not result in significant loss of information.

In the context of the recently proposed amyloid/tau/neurodegeneration (A/T/N) biomarker classification system^26^, our findings suggest that those characterized as A+/T-exhibit similar decline to those characterized as A+/T+ in late middle-age. However, we have not yet fully examined neurodegeneration. Total and phosphorylated tau were incorporated into the tau positivity classification and as they are highly correlated in this sample (*r* = .85, *p* < .001) it was not feasible to disambiguate neurodegeneration from neurofibrillary tau in this analysis. Furthermore, we did not observe differences among biomarker groups in hippocampal volume, unlike a prior study ^27^. It is possible these differences are due to the younger age of our cohort, which may not be expected to show structural brain changes at this stage, or that incorporation of additional structural imaging markers (e.g., cortical thickness) is needed to provide additional sensitivity and specificity to early neurodegeneration in AD.

Based on prior meta-analyses of cognitive decline in preclinical AD^14^ we focused on episodic memory and executive functioning measures; however, different patterns may be observed in other domains such as visuospatial function. It should be noted that factors that may be unrelated to AD can contribute to poor performance on cognitive tests (e.g., depression, sleep disorders, cerebrovascular disease) and continued longitudinal observation will be needed to parse the effects due to slowly evolving Aβ and tau pathology versus other explanations. Future analyses should examine additional differences between Aβ+ and Aβ− asymptomatic adults to determine if other factors (e.g., vascular risk factor burden) exacerbate decline in Aβ+ asymptomatic adults. An important limitation was inclusion of only CSF AD biomarkers and future analyses will incorporate CSF and molecular neuroimaging biomarkers to provide greater reliability in classification of preclinical AD. Our sample contained a smaller proportion of adults with markers of only tau+ (8%) compared to other studies (11-23%), perhaps due to the younger mean age of our cohort, the method by which we defined the cutoffs, or the relatively small sample from which the cutoffs were derived. These results are based on longitudinal cohorts that include a majority of Caucasian, highly educated adults from the Midwest region of the United States and may be less generalizable to other populations.

## ACKNOWLEDGMENTS

We gratefully acknowledge the assistance of researchers and staff at the Wisconsin Registry for Alzheimer’s Prevention and Wisconsin Alzheimer’s Disease Research Center for assistance in recruitment and data collection. Most importantly, we thank the dedicated WRAP and WADRC participants for their continued support and participation in this research.

